# Uncovering Long Palindromic Sequences in Rice (*Oryza sativa* subsp. *indica*) Genome

**DOI:** 10.1101/015099

**Authors:** Asadollah Ahmadikhah, Elmira Katanchi Kheiavi, Ali Mohammadian Mosammam

## Abstract

Because the rice genome has been sequenced entirely, search to find specific features at genome-wide scale is of high importance. Palindromic sequences are important DNA motifs involved in the regulation of different cellular processes and are a potential source of genetic instability. In order to search and study the long palindromic regions in the rice genome “R” statistical programming language was used. All palindromes, defined as identical inverted repeats with spacer DNA, could be analyzed and sorted according to their frequency, size, GC content, compact index etc. The results showed that the overall palindrome frequency was high in rice genome (nearly 51000 palindromes), with highest and lowest number of palindromes, respectively belongs to chromosome 1 and 12. Palindrome numbers could well explain the rice chromosome expansion (R^2^>92%). Average GC content of the palindromic sequences is 42.1%, indicating AT-richness and hence, the low-complexity of palindromic sequences. The results also showed different compact indices of palindromes in different chromosomes (43.2 per cM in chromosome 8 and 34.5 per cM in chromosome 3, as highest and lowest, respectively). The possible application of palindrome identification can be the use in the development of a molecular marker system facilitating some genetic studies such as evaluation of genetic variation and gene mapping and also serving as a useful tool in population structure analysis and genome evolution studies. Based on these results it can be concluded that the rice genome is rich in long palindromic sequences that triggered most variation during evolution.

**Availability***: The R scripts used to construct the palindrome sequence library are available upon request*.

## INTRODUCTION

Rice is one of the most important cereal crops around the world having the smallest genome among the cereals (Juretic et al. 2004). Rice is considered as a suitable model in genetic studies of cereal crops and other grasses. Repetitive sequences in rice genome are classified into transposable elements (TEs), sequences associated with centromere, telomeres, rRNA-related genes and other unknown sequences. Some of these repetitive sequences have known biological roles such as rRNA genes, centromeres and telomeres, while the role of other repetitive sequences stayed unknown (Yuan et al. 2003). Repetitive sequences significantly represent a candidates for regulatory sites (Horng et al. 2002). A main class of repetitive sequences are palindrome regions in genomes. Palindromes are present in both prokaryotes and eukaryotes and they involve in many biological processes, for example they act as recognition sites for activity of restriction enzymes, play an important role in DNA replication and regulation of gene expression through the formation of stem-loop structure (Ninh and Battig 2012).

Palindrome-containing sequences in mouse, yeast and bacteria genomes show a high instability. Inverted repeats of <10 bp distance cause B-DNA to form a cruciform structure (Rattray 2004). Short palindrome sequences (60–66 bp) named palindrome repeat sequences (PRS) were found in the rice genome. A single PRS was found in the intron region of mitochondrial gene *rps3* (ribosomal protein 3) (Nakazono and Kanno 1994). Study on distribution pattern of palindromes in yeast (*Saccaromyses cervisiea*) genome showed that long palindromes are rich in AT and mainly locate in intron regions of genes (Lisnić et al. 2005). A representative scheme showing the inverted repeat sequences in yeast is shown in Figure 1.

**Figure 1.**
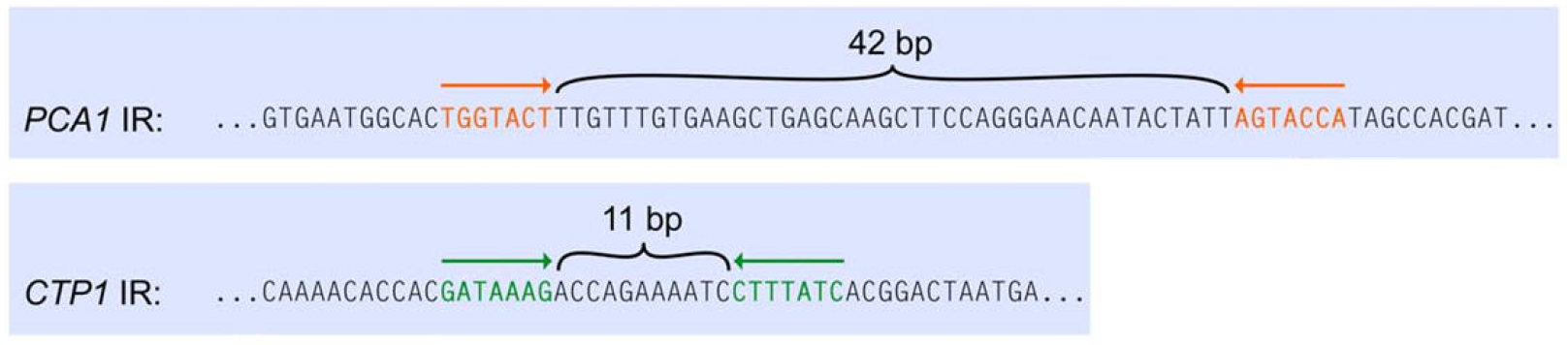
The inverted repeat sequences in *CTP1* and *PCA1* genes that define the breakpoints of a specific *SUL1* amplification event in yeast (Araya et al. 2010). Adopted from Brewer et al. (2011).

Genome studies showed that eukaryotic TE elements (transposons and retrotransposons) have short palindrome sequences or long palindrome sequences (named long terminal repeats, LTRs) in their structure. These sequences aren`t responsible for coding any protein, but instead they carry needed sequences for starting and terminating transcription of transposons. In grasses, LTR-retrotransposons can compose more than half of the genome of some species (San Miguel et al. 1996; Vicient et al. 1999; Kalendar et al. 2000; Schulman and Kalendar 2005). Majority of plant transposons are activated at stress condition. Thus, they can be desirable elements for the organism (Rashidi Monfared et al. 2009).

Pattern analysis of tandem repeats in genes is an indispensable computational approach to understand the gene expression and pathogenesis of diseases in humans (Tee 2013). The high mutation rate in tandem repeats is primarily due to the error in DNA replication that leads to the changes in the number of repeat units. In addition, it is also known that variable tandem repeats in the genome of an organism can accelerate the evolution of coding and regulatory gene sequences (Gemayel et al. 2010). The information obtained from the investigation of tandem repeats has widespread application in various fields such as medicine, forensic science, and population genetics (Xu et al. 1997). Notably, the identification of tandem repeats has special function in genetic studies, such as gene mapping (Dong et al. 2009). Tandem repeats are a stretch of nucleotides that repeat in a consecutive manner. They are ubiquitous in genome and their mutated forms are responsible for diseases (Gemayel et al. 2010). Understanding of the patterns of tandem repeats may provide insights into genome evolution and gene expression patterns. The advancement of computational tools such as computer algorithms, databases and web servers provide an efficient avenue to analyze enormous volume of genetic data, such as tandem repeats in large genome.

Location of these elements inside or near the active genes causes to start or end of the gene activity. Selection of new genes are produced looking for mutation or gene integration, requires forces that will determine the evolution of organisms (Ahmadikhah 2013; Yang et al. 2001; Rao 2010; Linheiro and Bergman 2012). Identification of repetitive regions provides insight into expression patterns of disease genes. Development of computational tools such as computer algorithms, databases and web servers, has made it possible to develop *in silico* approaches suitable for analysis of genetic data, including constituent repetitive sequences in the entire genome. R is both a language and an interface for statistical analysis, programming, and graphics. R is modeled after the S language that was originally created by AT&T and in many cases scripts written for R can be run in S with little to no modification. R has become a standard interface for statistical analysis in biological sciences due in part to its openness, ability to be extended by users and it vibrant user base. (Dyer 2009).

Regarding that the statistical programming language R has become a *de facto* standard for the analysis of many types of biological data (Wittelsbürger et al. 2015), we attempted to utilize this programming language for identifying long palindrome sequences in the rice (*O. sativa* subsp. *indica*) genome and for extracting some of specific features including GC content and nucleotide composition of the detected palindrome sequences in each rice chromosome.

## METHODS

### Data sets and searching for palindrome regions

At first step, the genomic sequences of 12 chromosomes of rice (*Oryza sativa* subsp. *indica*) were retrieved from NCBI website (www.ncbi.nlm.nih.gov) and stored in a PC drive in “FASTA” format to create early data set. FASTA files were called in R, and “Biostrings” package and “findPalindrome” command were used to search palindromic regions in each rice chromosome. The criteria for isolating palindromic regions were as follow: minimum arm length=20 nucleotides, minimum loop length=0 nucleotides (because existed available “findPalindrome” command doesn`t permit other numbers), maximum loop length=10000 nucleotides, and allowing no mismatch (maximum mismatch=0). A Perl programme was used to call the results of R programme. Secondary commands were written to filter out unwanted results such as very short palindromes (by setting minimum loop length=60 nucleotides). Final outputs were stored to create secondary data set.

### GC content

GC content of each palindrome region (partitioned in stem, loop and total) was calculated using “Biostrings” and “seqinr” packages.

### Compactness index

Frequency of palindromic sequences were separately calculated for each chromosome in one million intervals. Compactness index (CI) in view of the number of palindromes in each interval and in whole chromosome was calculated as the number of palindromes per Mbp and or per cM (given that 1 cM = 280 Kbp in rice genome).

## RESULTS

### Palindrome frequency

Searching for palindromic regions in the rice genome showed that rice genome has high frequency of palindromic regions (Supplementary File 1). In total, with our searching criteria, rice genome has nearly 51000 palindrome sequences, with an average of 4243 palindrome regions per chromosome (Table 1, Figure 2). Chromosome 1 has the highest (6598) and chromosome 12 has the least (2993) number of palindromes. Palindrome regions totally cover 41.4% of nuclear genome of rice, with highest coverage (52.9%) for chromosome 10 and lowest coverage (32.3%) for chromosome 3. For more details see Supplementary File 2.

**Table 1.** Number of palindrome regions in 12 *indica* rice chromosomes and some related statistics.

| Chromosome | Chromosome length (Kbp) | No. of palindrome regions | Chromosome coverage (%) | Average palindrome length (bp) | Average stem length (bp) | Average loop length (bp) |
| --- | --- | --- | --- | --- | --- | --- |
| 1 | 47244.9 | 6598 | 19524.7 (41.3%) | 2959.2(104-10792) | 33.9(20-1610) | 2891.4(61-9991) |
| 2 | 38080.4 | 4873 | 13234.4(34.8%) | 2715.9(101-10451) | 32.5(20-1585) | 2650.8(61-9977) |
| 3 | 41835.9 | 5159 | 13511.9(32.3%) | 2619.1(102-10099) | 32.7(20-867) | 2553.7(61-9988) |
| 4 | 34660.7 | 4341 | 12903.7(37.2%) | 2972.5(102-10988) | 33.4(20-1165) | 2905.6(61-10000) |
| 5 | 31162.6 | 4430 | 13882.6(44.5%) | 3133.8(101-10134) | 33.2(20-767) | 3068.4(61-9999) |
| 6 | 32845.4 | 4719 | 14631.1(44.5%) | 3100.5(102-10205) | 33.6(20-942) | 3033.3(62-9996) |
| 7 | 27898.3 | 3886 | 11584.3(41.5%) | 2981.0(101-10137) | 31.7(20-989) | 2917.6(61-9993) |
| 8 | 29367.1 | 4535 | 14496.0(49.4%) | 3196.5(102-10167) | 35.4(20-1104) | 3125.7(61-9993) |
| 9 | 21712.9 | 3031 | 9431.7(43.4%) | 3111.8(101-11574) | 34.1(20-1203) | 3043.5(61-9998) |
| 10 | 22150.1 | 3261 | 11726.9(52.9%) | 3596.1(102-10244) | 34.1(20-1418) | 3527.8(61-9990) |
| 11 | 22971.7 | 3088 | 9864.3(42.9%) | 3194.4(102-10259) | 33.0(20-1080) | 3128.4(61-9994) |
| 12 | 22948.0 | 2993 | 9450.9(41.2%) | 3157.7(102-11061) | 33.0(20-1337) | 3091.6(61-9979) |
| Total / mean | 372887.9 | 50914 | 154242.5(41.4%) | 3029.5(101-11574) | 33.4(20-1610) | 2962.8(61-10000) |

**Figure 2.**
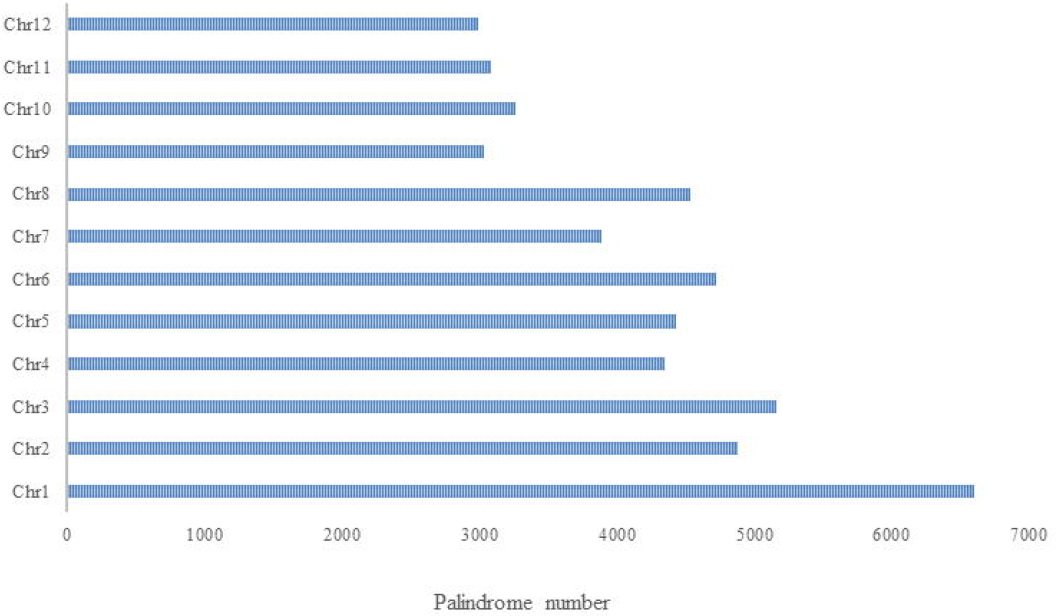
Distribution of palindrome regions in 12 rice chromosomes.

Average whole palindrome length is 3029 bp ranging from 101 to 11574 bp (Figure 3). The highest average palindrome length (3596 bp) belongs to chromosome 10 and the lowest (2619 bp) belongs to chromosome 3. Average stem length is 33.4 bp ranging from 20 to 1610 bp. The highest average stem length (35.4 bp) belongs to chromosome 8 and the lowest (31.7 bp) belongs to chromosome 7. Average loop length is nearly 2963 bp ranging from 61 to 10000 bp. The highest average loop length (3527 bp) belongs to chromosome 10 and the lowest (2553 bp) belongs to chromosome 3.

**Figure 3.**
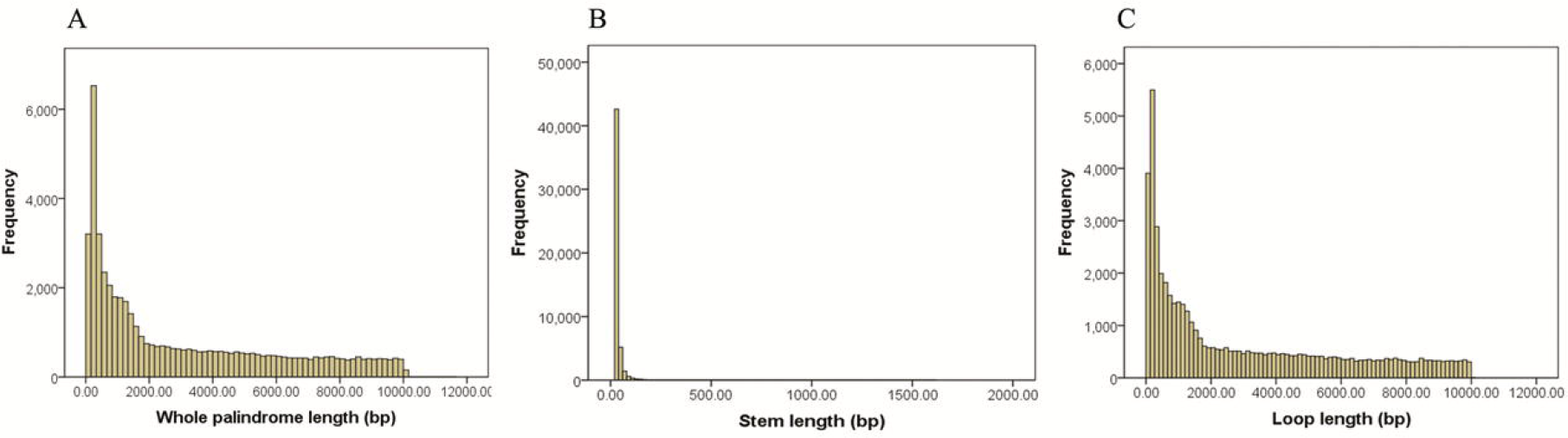
Frequency distribution of whole palindrome length (A), stem length (B) and loop length (C) for detected palindrome regions in rice nuclear genome (for more details see Supplementary File 3).

Correlation analysis showed that palindrome number has a high positive effect on chromosome expansion (R^2^>92.5%); in contrast palindrome length has a relatively high negative effect on chromosome expansion (R^2^>51%) that it can be explained by the size of loop section (and not by that of stem section) of palindrome regions (Figure 4).

**Figure 4.**
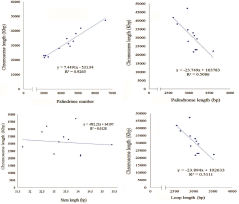
Effect of palindrome number, palindrome length, stem length and loop length on rice chromosome expansion.

### GC content

Average GC contents of palindrome regions in 12 chromosomes are depicted in Table 2. As seen, average of whole GC contents of the palindromes in rice genome is 42.1%, ranging from 41.0% (in chromosome 11) to 43.3% (in chromosome 5). The highest stem GC content (42.6%) was calculated for chromosome 10 and the lowest (40.0%) for chromosomes 11 and 12. The highest loop GC content (43.2%) was calculated for chromosome 5 and the lowest (40.8%) for chromosome 11. Based on these results, it seems that average GC contents of loops are slightly more than that of stems (42.0% vs. 40.9% in average). A graph showing stem and loop GC contents of palindrome regions of 12 chromosomes is presented in Figure 5.

**Table 2.** Average GC content of all palindrome regions for 12 *indica* rice chromosomes. Standard deviations are shown in parenthesis.

| Chromosome | Stems GC | Loops GC | Whole GC<br>(stems+loops) |
| --- | --- | --- | --- |
| 1 | 40.7(±16.9) | 42.2(±12.2) | 42.3(±11.6) |
| 2 | 40.3 (±16.4) | 42.0(±12.6) | 42.1(±11.9) |
| 3 | 40.9(±11.9) | 42.0(±16.9) | 42.2(±12.6) |
| 4 | 41.0(±11.4) | 41.6(±16.7) | 41.70(±12.0) |
| 5 | 41.2(±11.5) | 43.2(±16.6) | 43.3(±12.1) |
| 6 | 41.1(±11.8) | 42.6(±17.2) | 42.7(±12.4) |
| 7 | 41.3(±11.5) | 42.0(±17.7) | 42.1(±12.0) |
| 8 | 40.2(±11.0) | 41.5(16.4) | 41.6(±11.5) |
| 9 | 41.3(±11.2) | 41.5(±16.9) | 41.6(±11.8) |
| 10 | 42.6(±8.4) | 42.4(±16.1) | 42.6(±9.0) |
| 11 | 40.0(±11.2) | 40.8(±16.5) | 41.0(±11.9) |
| 12 | 40.0(±11.0) | 41.0(±16.1) | 41.1(±11.8) |
| Grand mean | 40.9(±16.8) | 42.0(±12.0) | 42.1(±11.3) |

**Figure 5.**
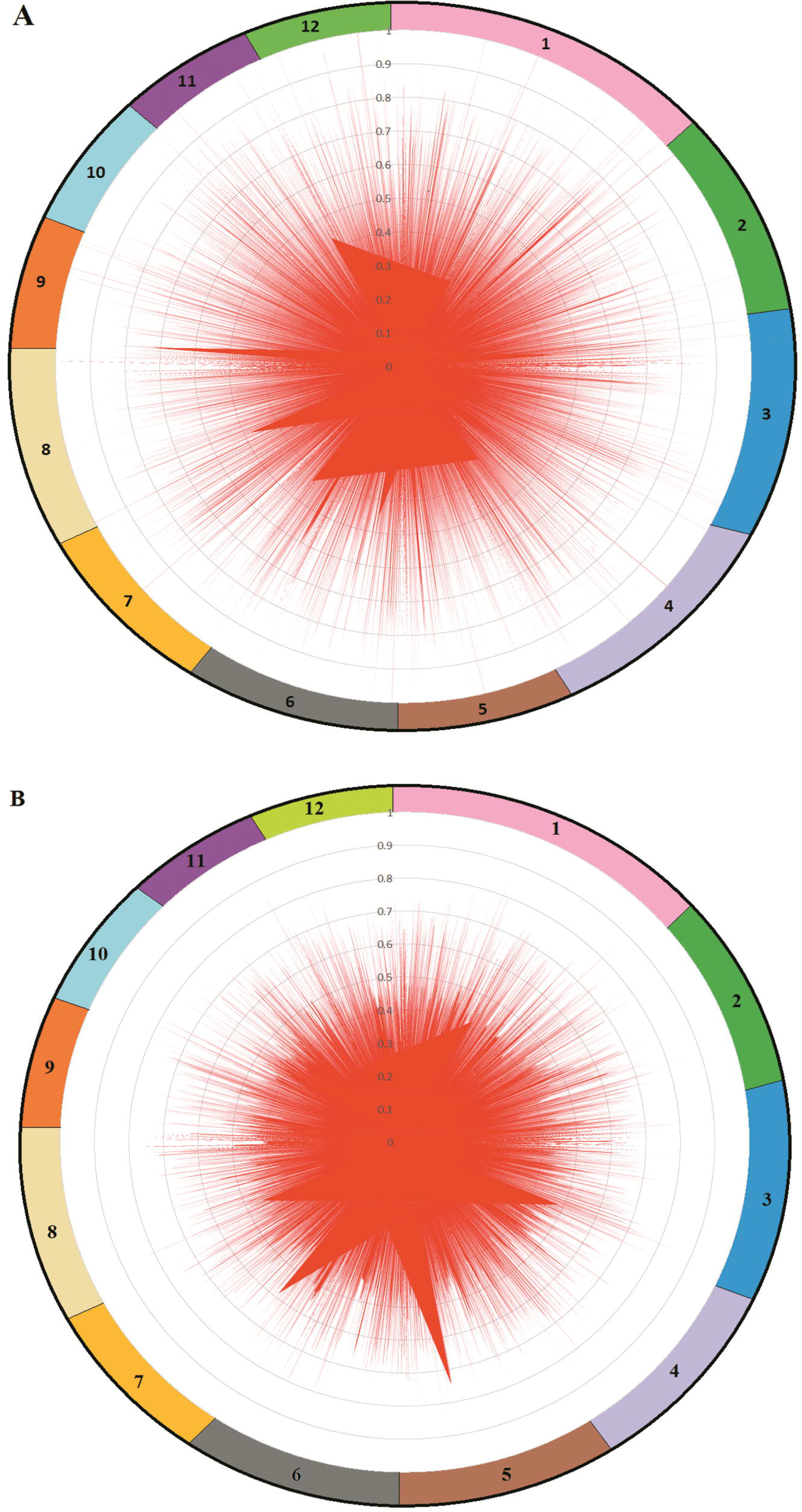
Frequency distribution of stem GC contents (A) and loop GC contents (B) of palindrome regions in 12 rice chromosomes (For more details see Supplementary File 4).

More detailed analysis on the abundance and frequency of GC-rich palindrome sequences (Table 3) showed that 6824 (13.4%) and 4316 (8.5%) of the detected palindromes (out of 50914) are GC-rich in stems and loops sections, respectively. In contrast, 26516 (52.1%) and 20087 (39.5%) of palindromes are AT-rich in stems and loops sections, respectively.

**Table 3.** Abundance and frequency of GC- and AT-rich palindrome sequences calculated separately for stems and loops sections

| Chromosome | Stems |  |  |  | Loops |  |  |  |
| --- | --- | --- | --- | --- | --- | --- | --- | --- |
|  | GC-rich |  | AT-Rich |  | GC-rich |  | AT-Rich |  |
|  | *n* | *f*(%) | *n* | *f*(%) | *n* | *f*(%) | *n* | *f*(%) |
| 1 | 933 | 14.14 | 3472 | 52.62 | 638 | 9.67 | 2521 | 38.21 |
| 2 | 577 | 11.84 | 2607 | 53.50 | 504 | 10.34 | 2065 | 42.38 |
| 3 | 713 | 13.82 | 2718 | 52.68 | 498 | 9.65 | 2097 | 40.65 |
| 4 | 581 | 13.39 | 2210 | 50.91 | 343 | 7.90 | 1674 | 38.56 |
| 5 | 588 | 13.27 | 2271 | 51.26 | 417 | 9.41 | 1540 | 34.76 |
| 6 | 662 | 14.03 | 2425 | 51.39 | 439 | 9.30 | 1725 | 36.55 |
| 7 | 610 | 15.70 | 2027 | 52.16 | 312 | 8.03 | 1529 | 39.35 |
| 8 | 556 | 12.26 | 2419 | 53.34 | 337 | 7.43 | 1908 | 42.07 |
| 9 | 425 | 14.02 | 1529 | 50.44 | 233 | 7.69 | 1204 | 39.72 |
| 10 | 492 | 15.09 | 1520 | 46.61 | 155 | 4.75 | 1172 | 35.94 |
| 11 | 366 | 11.85 | 1695 | 54.89 | 239 | 7.74 | 1414 | 45.79 |
| 12 | 312 | 10.42 | 1623 | 54.23 | 192 | 6.42 | 1238 | 41.36 |
| Total | 6824 | 13.40 | 26516 | 52.08 | 4316 | 8.48 | 20087 | 39.45 |
| n, number; f, frequency |  |  |  |  |  |  |  |  |

### Palindrome occurrence and compact indices in the rice genome

The results of compact indices (in view of palindromes/Mbp and palindromes/cM) are depicted in Table 4. In view of palindrome number per one Mbp, in average 136.6 palindromes occurred every one Mbp in the rice genome. Chromosome 8 has the highest (154.4±57.3) and chromosomes 3 has the lowest (123.3±39.6) palindrome numbers per Mbp. However, in view of palindrome occurrence per cM, in average 38.2 palindromes per cM exist in the rice genome (given that 1 cM contains 280 Kbp). However, different chromosomes have different compact indices, so that chromosome 8 has the highest CI (43.2 palindromes per cM), and chromosome 3 has the least CI (34.5 palindromes per cM), (Figure 6; Supplementary File 5).

**Table 4.** Compactness indices (CIs) and occurrence of palindromes in rice genome

| Chromosome | Length (cM) | Palindromes/Mbp | Palindromes/cM | Occurrence<br>(bp/palindrome) |
| --- | --- | --- | --- | --- |
| 1 | 168.73 | 139.7 ($\pm 54.9$) | 39.1 ($\pm 14.8$) | 7160.5 ($\pm 2612.6$) |
| 2 | 136.00 | 128.0 ($\pm 57.3$) | 35.8 ($\pm 16.0$) | 7814.6 ($\pm 3955.7$) |
| 3 | 149.41 | 123.3 ($\pm 39.6$) | 34.5 ($\pm 11.2$) | 8109.3 ($\pm 2974.7$) |
| 4 | 123.79 | 125.2 ($\pm 41.6$) | 35.1 ($\pm 11.4$) | 7984.5 ($\pm 3080.3$) |
| 5 | 111.29 | 142.2 ($\pm 58.5$) | 39.8 ($\pm 15.4$) | 7034.4 ($\pm 3035.8$) |
| 6 | 117.300 | 143.7 ($\pm 61.3$) | 40.2 ($\pm 17.0$) | 6960.2 ($\pm 3685.8$) |
| 7 | 99.63 | 139.3 ($\pm 60.4$) | 39.0 ($\pm 17.1$) | 7179.2 ($\pm 3221.8$) |
| 8 | 108.45 | 154.4 ($\pm 57.3$) | 43.2 ($\pm 19.7$) | 6475.7 ($\pm 3458.2$) |
| 9 | 77.54 | 139.6 ($\pm 51.3$) | 39.1 ($\pm 13.9$) | 7163.6 ($\pm 2886.9$) |
| 10 | 79.10 | 147.2 ($\pm 54.5$) | 41.2 ($\pm 13.9$) | 6792.4 ($\pm 3131.6$) |
| 11 | 82.04 | 134.4 ($\pm 38.4$) | 37.6 ($\pm 10.7$) | 7439.0 ($\pm 2008.3$) |
| 12 | 81.95 | 130.4 ($\pm 60.1$) | 36.5 ($\pm 16.8$) | 7667.2 ($\pm 2600.3$) |
| Mean | | 136.6 ($\pm 53.3$) | 38.2 ($\pm 15.1$) | 7323.9 ($\pm 3116.9$) |

**Figure 6.**
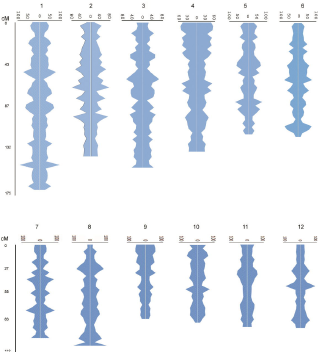
Distribution of palindrome compactness indices (CIs) along the 12 rice chromosomes. A vertical scale at left indicates the positions in cM and horizontal scales at the top of each chromosome indicate the CI values.

Assessment of the occurrence of palindrome regions in rice genome showed that in average one palindrome region occurs every 7323 bp (Table 4). However, the pattern of this occurrence is very variable between different chromosomes as well as in different regions of a specific chromosome (Figure 6). As seen in the figure, convexities represent a high compactness, indicating that repetitive sequences including palindrome regions play a vital role in heterochromatinization of the rice genome. Among rice chromosome segments, short arms of chromosomes 4 and 9 show a highest compactness. Almost for all rice chromosomes, centromeric regions show highest compactness (occurrence of higher palindromes per cM). In contrast, telomeric regions don`t follow a general rule of palindrome occurrence. As seen in Figure 6, telomeric side of short arms of chromosomes 3, 6, 8, 11 and 12, and telomeric side of long arms of chromosomes 2, 5, 6, 9, 10 and 12 have a low compactness (occurrence of lower palindromes per cM).

## DISCUSSION

In this research we succeeded to search and find many number of palindromic sequences in the rice (*O. sativa* subsp. *indica*) genome. As bioinformatics analyses showed rice genome is occupied by nearly 51000 long palindrome regions, and these regions occupies more than 41% of the rice nuclear genome. In a research it was reported that only 5% of vertebrate mitochondrial DNA has palindrome sequences, and in contrast mammalian mitochondrial genome hasn’t palindrome sequences (Arunkumar and Nagaraju 2006). Regarding that these types of sequences are classified as repetitive elements, a study showed that around 51% of the rice genome is occupied by repetitive elements (Horng et al. 2002). In other hand, transposable elements as a main class of repetitive sequences are virtually ubiquitous and make up, for instance, 20% of a typical *D. melanogaster* genome (Bergman et al. 2006), 50% of a *H. sapiens* genome (Lander et al. 2001), and 85% of a *Z. mays* genome (Schnable et al. 2009). Why repetitive sequences including transposons and palindromic regions have such an extended distribution in genomes, is probably related to their known or unknown functions in genome evolution. For example, repetitive extragenic palindromic sequences are DNA targets for insertion sequence elements in bacteria (Tobes and Pareja 2006). Palindromic sequences are known to have roles in DNA replication (Willwand et al. 1998) and RNA transcription (Chu et al. 1997). Also it has been reported that palindromes control gene expression through interaction with transcription factors, they maybe stabilize mRNA by inhibiting nuclease activity; they have been shown to involve in mRNA localization, and organisms use palindromes as markers for self-DNA and non-self-DNA; palindromes maybe prevent from the expression of foreign genes, and they are involved in correct intron splicing; furthermore, most palindromes act as recognition sites for DNA-binding proteins. (Giel-Pietraszuk et al. 2003; Tobes and Pareja 2006). Palindrome sequence plays a critical role in human foamy virus (HFV) dimerization (Cain et al. 2001). The analysis of repetitive elements also revealed that repetitive elements in human genome may have been very important in the evolutionary genomics (Horng et al. 2002).

Results showed that the distribution pattern of palindromes in rice chromosomes is adequately diverse, so that different chromosomes contain a different compact indices of palindromes. For example chromosome 3 has 123 palindromes per Mbp, indicating that the chromosome is a euchromatic chromosome governed a low diversity during evolution. In contrast, chromosome 8 has a high compact index of 154 palindromes per Mbp, most probably due to the high mobility of transposable elements in this chromosome (Bigot et al. 1990).

In general, we can classify rice palindrome regions into short (<100 bp) and long (>100 bp) palindromes. Results of our research showed that the detected palindrome regions have a different lengths, so that whole palindrome length ranged between 101 to 11574 bp (with an average of ∼3030 bp), and stem lengths ranged between 20 to 1610 bp (with an average of ∼33 bp), and loop lengths ranged between 61 to 10000 bp (with an average of 2963 bp). As before notified these long palindromes are over-represented in the rice nuclear genome (nearly 51000 long palindromes). Similarly, Lisnić et al. (2005) reported that while the short palindromes (2–12 bp) were under-represented, the palindromes longer than 12 bp were over-represented in the yeast genome. Apparently, palindrome length plays a critical role in genome stability. Palindromic sequences have been tied to different genomic rearrangements in different organisms depending on the length of the repeated sequences, although such repeated sequences are usually short and present at several functionally important regions in the genome. However, long palindromic sequences in yeast (*Saccharomyces cerevisiae*) genome are a major source of genomic instability (Nasar et al. 2000; Arunkumar and Nagaraju 2006; Sheari et al. 2008). The palindrome-mediated genomic instability is believed to be due to cruciform or hairpin formation and subsequent cleavage of this structure by structure-specific nucleases. Shorter palindromic sequences (shorter than 30 bp) are very stable while long palindromic sequences (>50 bp) generate double-strand breaks (DSBs) at a high frequency during meiosis in the yeast (*Saccharomyces cerevisiae*) that are not stable in vivo (Nasar et al. 2000). These sequences also increase inter- and intra-chromosomal recombination between homologous sequences. Hairpin structures can form from palindromic sequences due to base pairing in single-stranded DNA. These structures can be substrates for structure-specific nucleases and repair enzymes which can lead to a double-strand break (DSB) in the DNA. This then leads to loss of genomic material which can cause meiotic recombination (Nasar et al. 2000; Arunkumar and Nagaraju 2006; Sheari et al. 2008). Studies with a 140-bp long mutated palindromic sequence inserted in yeast have shown to lower post-meiotic segregation and increase rate of gene conversions, while shorter sequences do the opposite (Nag and Kurst 1997).

Correlation analysis showed that palindrome number plays a significant role in the expansion of chromosome size (R^2^>92%; Figure 4), although palindrome length has a relatively high negative effect on chromosome size; that is, the longer palindromes can be found in shorter chromosomes. This reverse effect causes a balanced occupation rate in different rice chromosomes, so that except for chromosomes 2 and 3, more than 40% of chromosome lengths are occupied by palindromic sequences (see Table 1). To verify this hypothesis, we calculated a more sensible index named PL:CL ratio (Figure 7). To do this, we divided average palindrome length (PL) of each chromosome by the respective chromosome length (CL). As seen in the figure, this ratio well explains (R^2^>90%) the negative effect of palindrome length on chromosome expansion. Piegu et al. (2006) concluded that accumulation of repetitive elements, particularly retrotransposable elements, besides polyploidy, is the main factor of genome size increase in higher eukaryotes. They showed that the genome of *Oryza australiensis*, a wild relative of the Asian cultivated rice (*O. sativa*), has accumulated more than 90,000 retrotransposon copies during the last three million years, leading to a rapid twofold increase of its size.

**Figure 7.**
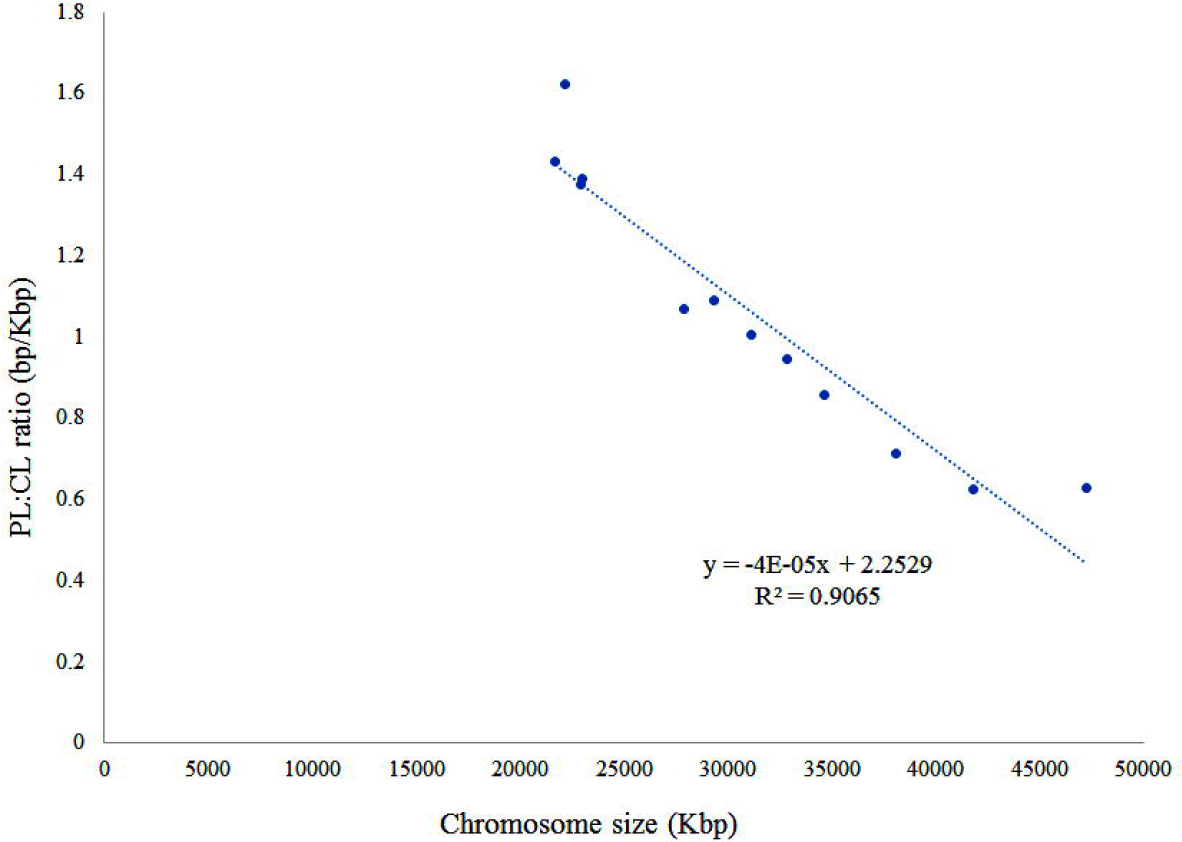
Relationship of PL:CL ratio on chromosome size in rice. PL, palindrome length; CL, chromosome length.

Results of our research also showed that one palindrome sequence occur every 7.3 Kbp in the rice genome, although this characteristics differed between chromosomes and even between the genomic locations. This value is considerably lower than that of (18.8 Kbp) reported for simple sequence repeats (SSR) in the rice genome (Li et al. 2004). These findings show that rice genome is rich in the palindrome sequences. A simulation study showed that there are 100 palindromes in every 1000 base pairs of a randomly generated sequence (Ninh and Battig 2012).

GC content analysis showed that average GC content of all detected whole-length palindromes is 42.1%, with being 40.9% and 42.0% the GC contents of stems and loops, respectively, indicating that the long palindrome sequences are GC-poor, although a considerable number of palindromes doesn’t follow this general rule (Figure 5). In fact, 13.4% and 8.5% of palindromes are GC-rich in stems and loops sections, respectively. In contrast, 52.1% and 39.5% of palindromes are AT-rich in stems and loops sections, respectively (Table 3). These results show the low-complexity of palindrome regions in the rice genome. Similarly, Sheari et al. (2008) reported that there was a large chance of finding a palindrome in low complexity sequences. Many low-complexity regions are highly unstable due to the action of replication slippage and recombination (Ellegren 2004), and the uncontrolled expansion of short sequence motifs causes several human diseases, such as Huntington’s disease and other neurodegenerative disorders (Gatchel and Zoghbi 2005). A hypothesis is that these regions increase phenotypic variation within populations, facilitating adaptation (Kashi and King 2006). An alternative hypothesis to explain the abundance of low-complexity regions is that they facilitate the formation of novel coding sequences (Green and Wang 1994; Toll-Riera et al. 2012). In addition to the possible application of data obtained by palindrome identification in the development of a marker system for the use in some genetic studies such as evaluation of genetic variation and gene mapping (Ahmadikhah 2009; Dong et al. 2009), these findings can be serve as a useful tool in population structure analysis and genome evolution studies.

## CONCLUSION

Based on the results of this research it can be concluded that the rice genome is rich in long palindromic sequences that triggered most variation during evolution. Generally, both sections of palindromic sequences including stems and loops are AT-rich, indicating that these regions locate in the low-complexity segments of the rice chromosomes. The palindrome identification can help to develop a molecular marker system facilitating some genetic studies such as evaluation of genetic variation and gene mapping and also serves as a useful tool in population structure analysis and genome evolution studies.

## ACKNOWLEDGMENTS

We thank Shahid Beheshti University for annual grants and University of Zanan for partial financial support of the work.

